# The Reconstruction of 2,631 Draft Metagenome-Assembled Genomes from the Global Oceans

**DOI:** 10.1101/162503

**Authors:** Benjamin J. Tully, Elaina D. Graham, John F. Heidelberg

## Abstract

Microorganisms play a crucial role in mediating global biogeochemical cycles in the marine environment. By reconstructing the genomes of environmental organisms through metagenomics, researchers are able to study the metabolic potential of Bacteria and Archaea that are resistant to isolation in the laboratory. Utilizing the large metagenomic dataset generated from 234 samples collected during the *Tara Oceans* circumnavigation expedition, we were able to assemble 102 billion paired-end reads into 562 million contigs, which in turn were co-assembled and consolidated in to 7.2 million contigs ≥2kb in length. Approximately 1 million of these contigs were binned to reconstruct draft genomes. In total, 2,631 draft genomes with an estimated completion of ≥50% were generated (1,491 draft genomes >70% complete; 603 high-quality genomes >90% complete). A majority of the draft genomes were manually assigned phylogeny based on sets of concatenated phylogenetic marker genes and/or 16S rRNA gene sequences. The draft genomes are now publically available for the research community at-large.

## Background & Summary

The global oceans are a vast environment in which many key biogeochemical cycles are performed by microorganisms, specifically the Bacteria and Archaea^1,2^. Assessing the role of individual microorganisms has been confounded due to limitations in growing and maintaining ‘wild’ organisms in the laboratory environment^3^. The advent of “-omic” techniques, metagenomics, metatranscriptomics, metaproteomics, and metabolomics, has provided an avenue for exploring microbial diversity and function by skipping the necessity of culturing organisms, thus allowing researchers to study organisms for which growth conditions cannot be replicated.Specifically, the application of metagenomics, the sampling and sequencing of genetic material directly from environment, provides an avenue for reconstructing the genomic sequences of environmental Bacteria and Archaea^4-6^.

Through the *Tara Oceans* Expedition (2003-2010), thousands of samples were collected of marine life^7^, including more than 200 metagenomic samples targeting the viral and microbial components of the marine ecosystem from around the globe^8,9^. Several studies have started the process of reconstructing microbial genomes from these metagenomics samples, utilizing samples from the Mediterranean^10^ and the bacterial size fraction (0.2-3μm)^11^. Here, we present >2,000 additional draft genomes from the Bacteria and Archaea estimated to be >50% complete reconstructed from 102 billion metagenomic sequences generated from multiple size fractions and depths at the 61 stations sampled during the *Tara Oceans* circumnavigation of the globe.Phylogenomic analysis suggests that this set of draft genomes includes highly sought after genomes that lack cultured representatives, such as: Group II (149) and Group III (12) Euryarchaeota, the Candidate Phyla Radiation (30), the SAR324 (18), the *Pelagibacteraceae* (32), and the *Marinimicrobia* (111).

We envision that these draft genomes will provide a resource for downstream analysis acting as references for metatranscriptomic^12^ and metaproteomic^13^ projects, providing the data necessary for large-scale comparative genomics within globally vital phylogenetic groups^14^, and allowing for the exploration of novel microbial metabolisms^15^. Non-redundant draft metagenome-assembled genomes have been deposited into the National Center for Biotechnology Information (NCBI) database, along with publically accessible datasets for examining metagenomic information that was not incorporated in to the draft genomes.

## Methods

These methods have been described in part previously^15^, but have not been applied to full dataset discussed below.

### Gathering Metagenomics Sequences & Assembly

An example of the methodology used to assemble the *Tara Oceans* metagenomes is available on Protocols.io (dx.doi.org/10.17504/protocols.io.hfqb3mw). All metagenomic sequences generated for 234 samples collected from 61 stations during the *Tara Oceans* expedition were accessed from the European Molecular Biology Laboratory-European Bioinformatics Institute (EMBL-EBI)^8,9^. Generally, samples were collected from multiple size fractions, commonly ‘viral’(<0.22μm),‘girus’ (0.22-0.8μm), ‘bacterial’ (0.22-1.6μm), and ‘protistan’ (0.8-5.0μm), at multiple depths, commonly at the surface (~5-m), deep chlorophyll maximum (DCM), and mesopelagic, from each station. Each sample was assembled individually using Megahit^16^ (v.1.0.3; parameters: --preset meta-sensitive). In total, over 102 billion paired-end reads were assembled into >562 million contigs (Table 1; referred to as primary contigs).Primary contigs <2kb in length were not used in downstream analysis. Primary contigs ≥2kb in length were processed using CD-HIT-EST^17^ (v4.6; parameter: -c 0.99) to reduce the computational load required for the secondary assembly by combining contigs with ≥99% semi-global identity. Primary contigs for stations from the same oceanographic province were co-assembled using Minimus2^18^ (Figure 1; AMOS v3.1.0; parameters: -D OVERLAP=100 MINID=95). Combining the Minimus2 generated contigs and the primary contigs that did not assemble with Minimus2, approximately 7.2 million contigs were generated for downstream analysis (Table 2; referred to as secondary contigs).

**Figure 1.**
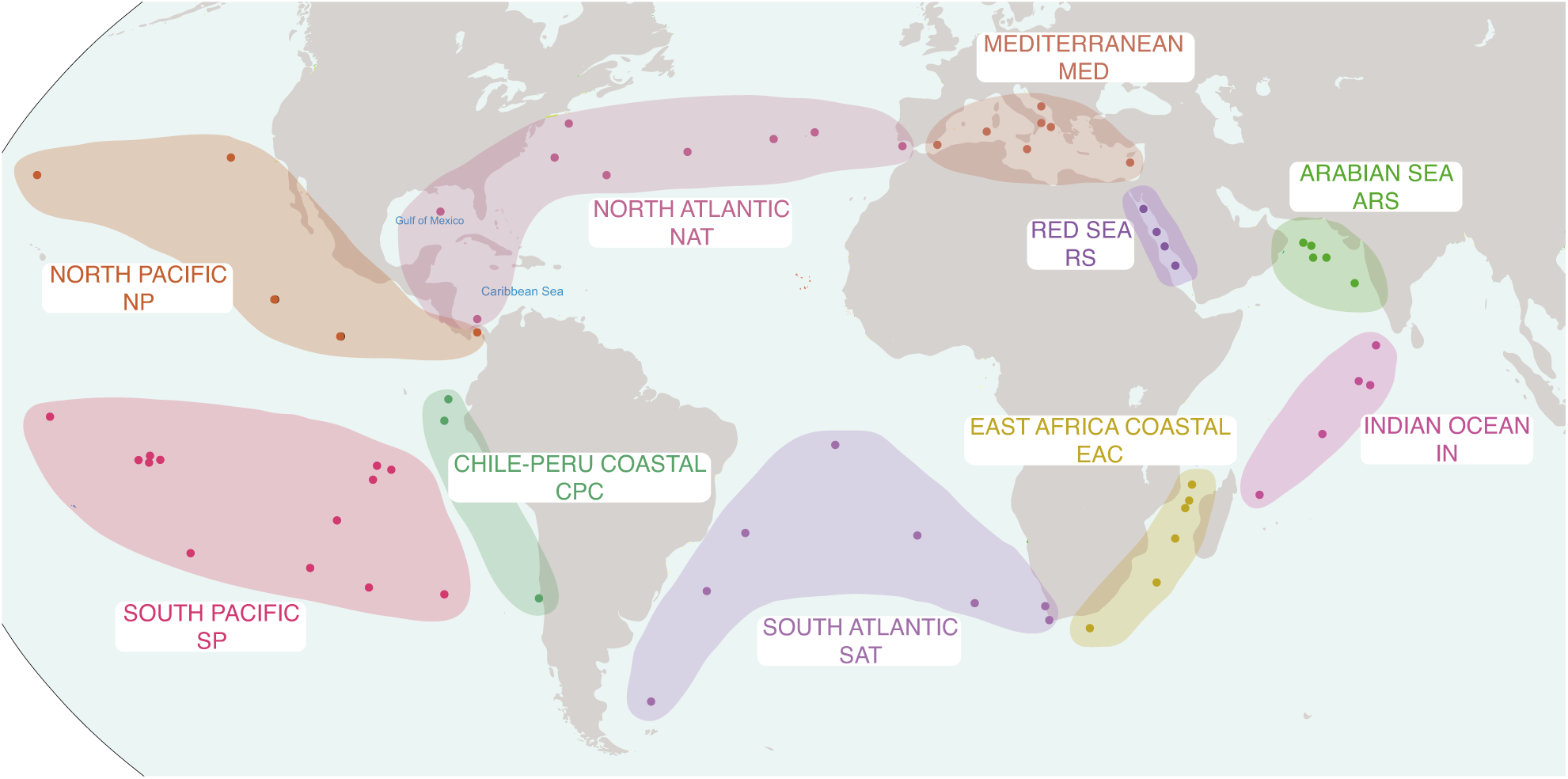
A map depicting the approximate locations of the *Tara Oceans* sampling stations from which metagenomics data was collected. Stations are grouped in to larger provinces based on Longhurst Provinces and site proximity. Province abbreviations are used for draft genome IDs. The map in Figure 1 were modified under a CC BY-SA 3.0 license from ‘Oceans and Seas boundaries map’ by Pinpin.

**Table 2.** Statistics for each province on the number secondary contigs generated, the number of contigs binned and corresponding length cutoff, and the number of draft genomes reconstructed.

| Province | No. of Secondary Contigs | Size Cutoff (kb) | No. of Binned Contigs | No. of Draft Genomes |
| --- | --- | --- | --- | --- |
| Mediterranean | 660,937 | 7.5 | 95,506 | 360 |
| Red Sea | 328,325 | 5.0 | 84,936 | 180 |
| Arabian Sea | 525,636 | 6.0 | 99,649 | 194 |
| Indian Monsoon | 285,238 | 4.0 | 93,760 | 72 |
| East Africa Coastal Current | 613,778 | 7.0 | 91,053 | 208 |
| South Atlantic | 1,373,173 | 11.5 | 96,972 | 360 |
| Chile Peru Coastal | 857,548 | 5.5 | 95,557 | 146 |
| South Pacific | 807,193 | 14.0 | 104,598 | 536 |
| North Pacific | 943,809 | 7.0 | 96,396 | 254 |
| North Atlantic | 804,316 | 8.5 | 104,848 | 321 |
| SUM | 7,199,953 | - | 963,275 | 2,631 |

### Binning

An example of methodology used to bin the *Tara Oceans* metagenomes is available on Protocols.io (dx.doi.org/10.17504/protocols.io.iwgcfbw). Metagenomic reads from each sample in a province were recruited against the set of secondary contigs generated for that province using Bowtie2^19^ (v4.1.2; default parameters). Utilizing the BinSanity^20^ workflow^21^ (BinSanity-profile), a reads·bp^-1^ coverage value was generated for each contig and coverage values were multiplied by 100 and log normalized (parameter: -transform scale). Then due to computational limitations imposed during the BinSanity binning method, the secondary contigs from each province were size selected (4-14kb cutoffs) to choose approximately 100,000 contigs for binning (Table 2). Approximately 6 million secondary contigs remain un-binned and are available for analysis. The binning using BinSanity was performed iteratively six times, with changes to the preference value after the first three iterations and a set parameter for iterations 4-6 in order to influence the degree of clustering (v0.2.5.5; parameters: -p [(1) -10, (2) -5, (3) -3,(4-6) -3] -m 4000 -v 400 -d 0.95 -kmer 4). Bins generated during the first five iterations were processed with the BinSanity-refinement script utilizing a set preference value (parameter: -p -25). After the six iteration, bins with high contamination (>10% contamination; see below) were processed two more times with BinSanity-refinement using variable preference values (parameter: -p [(6) -10, (7) -3]). After each refinement step, bins were assessed using CheckM^22^ (v1.0.3; parameters: lineage_wf) for completion and contamination estimates, which were used as cutoffs for inclusion in the final dataset. Bins were reassigned as a draft genome if: >90% complete with <10% contamination, 80-90% complete with <5% contamination, or 50-80% complete with <2% contamination. Bins that did not meet these criteria were combined for the next iteration of binning, except after the six iteration (see above). In total, 2,631 draft genomes were generated, with 1,491 of the genomes >70% complete, and 420 genomes meeting a high-quality threshold of >90% complete and <5% contamination (Table 3). Genomes were provided identifiers with the format ***T****ara* ***O****ceans* **B**inned **G**enome (TOBG) – Province Abbreviation – Numeric ID (*e.g.*, TOBG_NAT-221).

An additional 15,557 bins were generated containing at least five contigs that did not meet the criteria for reclassification as a draft genome. These bins may offer pertinent information for different downstream analyses. Bins of interest with high completion and high contamination can be manually assessed using tools, such as Anvi’o^23^, to generate a more accurate draft genome. For bins with <50% completion, it may be possible to combine two or more bins to generate a draft genome. And for bins with minimal or no phylogenetic markers assessment may reveal that they represent viral, episomal, or eukaryotic DNA sequences.

### Phylogenetic Assignment

A multi-pronged approach was used to provide a phylogenetic assignment to all of the draft genomes. All of the secondary contigs had putative coding DNA sequences (CDSs) predicted using Prodigal^24^ (v2.6.2; -m -p meta). Contigs assigned to draft genomes and 7,041 complete and partial reference genomes (Supplemental Table 1) accessed from NCBI GenBank^25^ and searched for phylogenetic markers. Protein phylogenetic markers were detected using hidden Markov models (HMMs) collected from the Pfam database^26^ (Accessed March 2017) and identified using HMMER^27^ (v3.1b2; parameters: hmmsearch -E 1e-10). Two sets of single-copy markers recalcitrant to horizontal gene transfer were identified and used to construct phylogenetic trees; a set of 16 generally syntenic markers identified in Hug^28^, *et al*. (2016) and an alternative set of 25 markers (Supplemental Table 2). Draft and reference genomes were required to possess ≥10 and ≥15 markers for the Hug, *et al*. and alternative marker sets, respectively, to be included in downstream analysis. If multiple copies of the same marker were detected, neither copy was considered for further analysis. Each marker was aligned using MUSCLE^29^ (v3.8.31; parameter: -maxiters 8), trimmed using trimAL^30^ (v.1.2rev59; parameter: -automated1), and manually assessed. Alignments for each set of markers were concatenated. A maximum likelihood tree using the LGGAMMA model was generated using FastTree^31^ (v.2.1.10; parameters: -lg -gamma; Supplemental Information 1 and 2). Phylogenies were determined manually for 2,009 and 95 draft genomes for the Hug, *et al*. and alternative marker sets, respectively (Table 4). A simplified phylogenetic tree of the Hug, *et al*. phylogenetic marker set was constructed using the same parameters with only the alignments of the draft genomes for Fig. 2.

**Figure 2.**
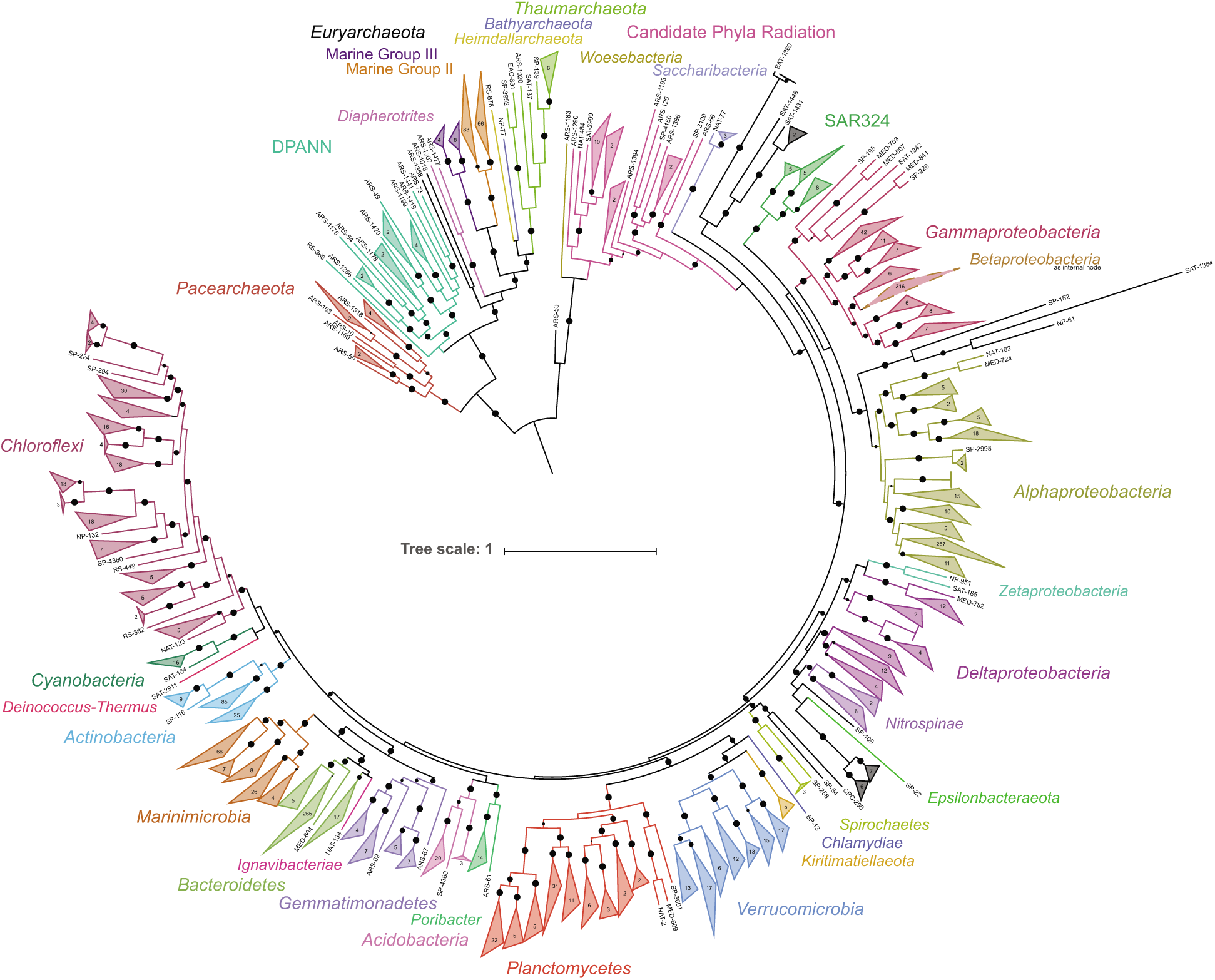
A maximum likelihood tree of the TOBG draft genomes based on 16 concatenated single-copy phylogenetic markers. Bootstrap values >0.75 are shown. Circle size representing the bootstrap value is scaled from 0.75-1.0. Nodes where the average branch length distance is <0.5 were collapsed and the number of draft genomes in each node are provided. The image was generated using the Interactive Tree of Life (iTOL; http://itol.embl.de/).

16S rRNA genes were predicted from draft genomes using RNAmmer^32^ (v1.2; parameters: -S bac -m ssu). 276 16S rRNA genes were detected and aligned using the SINA web portal aligner^33^ (https://www.arb-silva.de/aligner/). Aligned 16S rRNA gene sequences were added to the non-redundant 16S rRNA gene database (SSURef128 NR99) in ARB^34^ (v6.0.3) using the Parsimony (Quick) tool (default parameters). Each 16S rRNA gene sequence from a draft genome was assigned a putative phylogeny based on placement on the SSURef128 NR99 guide tree (Table 4).

For the draft genomes, 81.3% were manually assigned a phylogeny based on the Hug, *et al*. marker set (2,009 draft genomes), the alternative marker set (95 draft genomes), or the 16S rRNA gene tree (35 draft genomes). The remaining 492 draft genomes were provided a putative phylogeny based on CheckM (Table 4).

### Relative Abundance

Several of the size fractions used to reconstruct bacterial and archaeal draft genomes were specifically designed to target different biological entities, such as double-stranded DNA viruses, giant viruses (giruses), and protists. In order to estimate the relative abundance of the draft genomes compared to only the total bacterial and archaeal community, a set of 100 previously identified HMMs for predominantly single-copy bacterial and archaeal markers^35,36^ were searched against the putative CDS of the secondary contigs from each province using HMMER (parameters: hmmsearch --cut_tc). From each province, the set of CDS identified by the marker HMMs could be used to approximate the total bacterial and archaeal community. Markers belonging to the draft genomes were identified. Based on the metagenomic reads recruited to the secondary contigs for each sample, the number of reads aligned to each marker in a sample was determined using BEDTools^37^ (v2.17.0; multicov default parameters). A length-normalized estimate of relative abundance for each draft genome in each sample in a province was determined using the following equation:

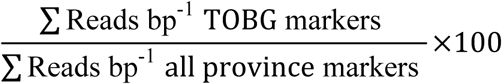

The relative abundance estimates of draft genomes indicate that the genomes generated for this study constitute only a small percentage of the total bacterial and archaeal abundance in each sample (Table 5; Figure 3). The draft genomes account for a higher percentage of the viral size fraction compared to other size fractions, accounting for ~60% of the total bacterial and archaeal community in that size fraction. This is likely due to the fact that the number of microbial organisms capable of passing through a 0.22μm filter is limited and the overall microbial community is these samples is less complex. On average, the draft genomes in the girus, bacterial, and protistan size fractions account for 14-19% of the total bacterial and archaeal communities. As such, the application of alternative binning methods to this same dataset should generate additional draft genomes^38^.

**Figure 3.**
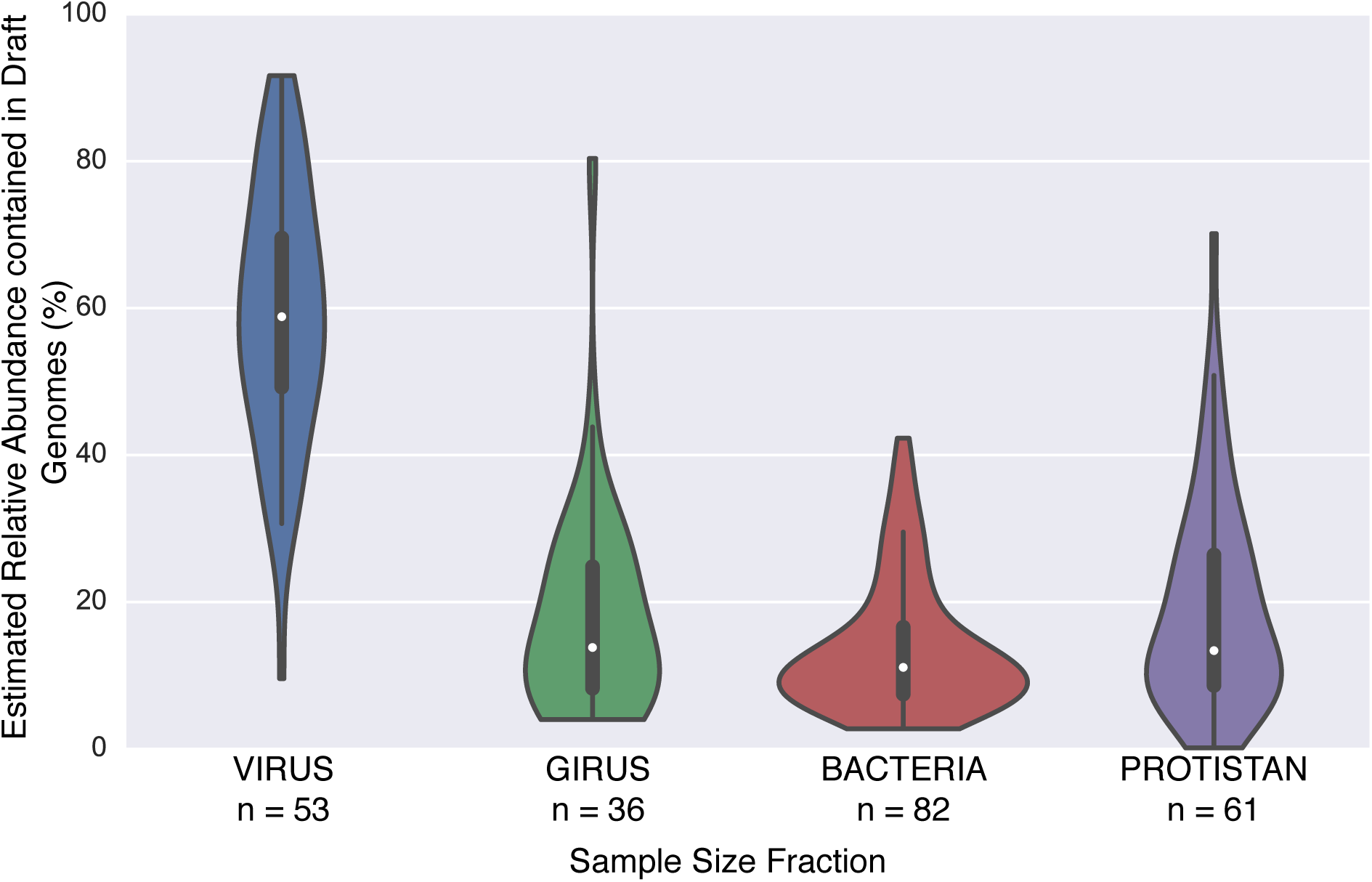
Violin plots illustrating the fraction of the estimated total bacterial and archaeal community represented by the draft genomes for samples from the different size fractions.

### Data Records

This project has been deposited at DDBJ/ENA/GenBank under the BioProject accession no. PRJNA391943 and drafts of genomes are available with accession no. XXX-XXX [submission to NCBI is ongoing – draft genomes can be found on provided figshare link] (Data Citation 1). Additional data is available through figshare, including all draft genomes, all secondary contigs, read count data for each secondary contig from each sample (Data Citation 2). The set of 100 HMMs for predominantly single-copy bacterial and archaeal markers from Albertsen, *et al*. (2013) is available on GitHub (Data Citation 3).

### Data Usage

Due to the draft nature of the TOBG genomes, all downstream research should independently assess the accuracy of genes, contigs, and phylogenetic assignments for organisms of interest. Several of the draft genomes generated through this methodology appear to be identical, based on the Hug marker set phylogenomic tree, to genomes generated by Tully, *et al*. (2017) and Delmont, *et al*. (2017), these genomes have been identified (Table 3) and in most cases duplicate genomes were not submitted to NCBI. In total, 186 draft genomes from this dataset, 68 from Tully, *et al*. (2017) and 118 from Delmont, *et al*. (2017), were determined to be identical to the previous work and not submitted to NCBI. However, draft genomes from this study that were estimated to be more complete than available through Delmont, *et al*. (2017) were submitted (n = 198) to NCBI. In providing official nomenclature for submission to NCBI, priority was given to the Hug marker assignment, followed by the 16S rRNA assignment, then alternative marker assignment, and, finally, the CheckM assignment.

### Tables

**Table 1.** Statistics for the primary contigs generated for each of the 234 sample fractions. (Due to size limitations in a bioRxiv submission, this table is included on the figshare page)

**Table 3.** Statistics for each of the 2,631 draft genomes, including completion and contamination. (Due to size limitations in a bioRxiv submission, this table is included on the figshare page)

**Table 4.** Phylogenetic assignment for each of the draft genomes as determined by the four methodologies outlined in the manuscript (Assignments for the Hug *et al.* marker gene set, alternative marker gene set, 16S rRNA gene, and CheckM). (Due to size limitations in a bioRxiv submission, this table is included on the figshare page)

**Table 5.** Estimated relative abundance value for all draft genomes in each sample fraction from each province. (Due to size limitations in a bioRxiv submission, this table is included on the figshare page)

## Data Citations

1. Tully, B. J. *NCBI BioProject* PRJNA391943 (2017)

2. Tully, B.J. *figshare* http://dx.doi.org/10.6084/m9.figshare.5188273 (2017)

*3.Albertsen, M. GitHub* https://github.com/MadsAlbertsen/multi-metagenome/blob/master/R.data.generation/essential.hmm (2011)

## Author Contribution

BJT conceived and designed the methodology, performed the analysis, wrote the paper, and prepared the figure and tables. EDG performed the analysis and reviewed drafts of the paper. JHF provided funding and resources to perform the analysis and reviewed drafts of the paper.

## Acknowledgements

Funding was provided by the Center for Dark Energy Biosphere Investigations (C-DEBI) to BJT and JFH (OCE-0939654). As we have sstated before, this project would have not been possible if not for the diligent commitment by the *Tara Oceans* consortium to allow for the open access of the data collected during the expedition. We only hope that this small dataset can be used by the scientific community at-large to increase the impact of this transformational research project. This is C-DEBI Contribution XXX.

